# Attention network modulation via tRNS correlates with attention gain

**DOI:** 10.1101/2020.09.15.298240

**Authors:** F. Contò, G. Edwards, S. Tyler, D. Parrott, E.D. Grossman, L. Battelli

## Abstract

Transcranial Random Noise Stimulation (tRNS) can enhance vision in the healthy and diseased brain. Yet, the impact of tRNS on large-scale cortical networks is still unknown. We investigated the impact of tRNS coupled with behavioral training on resting-state functional connectivity and attention. We trained human subjects for four consecutive days on two attention tasks, while receiving tRNS over the intraparietal sulci, the middle temporal areas, or sham stimulation. We measured resting state functional connectivity of nodes of the dorsal and ventral attention network (DVAN) before and after training. We found a strong behavioral improvement and increased connectivity within the DVAN after parietal stimulation only. Crucially, behavioral improvement positively correlated with connectivity measures. We conclude changes in connectivity is a marker for the enduring effect of tRNS upon behavior. Our results suggest that tRNS has strong potential to augment cognitive capacity in healthy individuals and promote recovery in the neurological population.

## Introduction

Vision and attention are the primary sensory and cognitive modalities through which humans interact with the environment. Augmentation of visuoperceptual function through training, termed perceptual learning (PL), is crucial for clinical and aging populations that benefit from therapeutic interventions to recover and retain perceptual function. PL is also a pivotal tool for investigating neural plasticity (Gilbert, 1994; Polat et al., 2004; Shibata et al., 2016; Deveau et al., 2014; Sterkin et al., 2018), and has been linked to localized neural changes in sensory cortex and long-range sensory-attention network reorganization (Gilbert et al., 2001; Li et al., 2004; Sasaki et al., 2010; Fahle, 2004). Thus, PL has potential as an approach to reveal underlying cortical reorganization that promotes sustained improvements in perceptual function.

Although promising, PL approaches have been limited in their applicability, due in part to the long and intensive training protocols which largely impede their use (Dosher and Lu, 2017; Das et al., 2014) (Li et al., 2004; Huang et al., 2012; Zhang et al., 2010a; 2010b; Dosher & Lu, 2005). Currently lacking are PL approaches to increase the rate of learning, valuable for populations with limited abilities to engage in demanding perceptual tasks. A notable exception is “cross-tasks” training, in which learning protocols are significantly shortened by adopting a training procedure on two different tasks with the same stimuli simultaneously (Szpiro, Wright and Carrasco, 2014, Wright et al., 2010).

More recently transcranial electrical stimulation (tES) has been proposed as a tool to facilitate cortical plasticity, particularly when coupled with behavioral training (Edwards et al., 2019). When delivered using a random noise protocol (tRNS), tES can modulate visual cortex excitability (Herpich et al., 2018), promote visual functions in healthy subjects (Fertonani et al., 2011; Pirulli et al., 2013; Tyler et al., 2018), and facilitate recovery of visual dysfunctions in disease (Campana et al., 2014; Camilleri et al., 2016; Herpich et al., 2019). Moreover, recent studies show PL coupled with transcranial electrical stimulation (tES) can successfully boost the effects of training by increasing the rate of learning across sessions (Fertonani et al., 2011), particularly when applied during task training (Pirulli et al., 2013), with the subsequent observed behavioral improvements persisting for durations well beyond those obtained through PL training alone (Cappelletti et al., 2013; Herpich et al., 2019).

Studies on the physiological effects of tRNS on brain dynamics are not yet clear on the mechanisms by which training is facilitated. The leading hypothesis derived from animal studies is that tRNS has a stochastic resonance effect upon neurons such that the injected subthreshold stimulation both resonates with ongoing task-related oscillatory activity and reduces endogenous noise, and thus, when applied at optimal levels, enhances the signal-to-noise of task-related spike firing (Antal and Herrmann, 2016; Liu et al. 2018; Polanía et al. 2018). Consistent with the network oscillation component of resonant theory, extant studies on humans show that focal tES of a node within a network can cause a cascade of functional changes selectively spreading within that network (Nitsche and Paulus, 2000; Polania et al., 2012; Antal et al., 2011; Krause et al., 2017). Modeling studies have likewise shown changes in network dynamics within the directly stimulated area and across distal but connected cortical areas (Kunze et al., 2015; Polania et al., 2011; Sehm et al., 2012). Given that many behavioral manifestations of neurological and psychiatric disorders are the consequence of altered brain network connectivity (Buckner, 2005; Fox et al., 2014), the network-wide impact of tES could have wide clinical applications.

The present study uses a combination of tRNS and fMRI, in conjunction with cross-training on a perceptual learning task, to investigate large-scale cortical dynamics when coupled with behavioral training. Specifically, we implemented a four-day attention training protocol coupled with tRNS to examine if tRNS to the parietal lobe would modulate the dorsal and ventral attention network, and subsequently increase attention. The target of the parietal stimulation was the intraparietal sulcus (IPS), a crucial node of spatial attention (Vossel, Geng and Fink, 2014; Battelli et al., 2017; Plow et al., 2014; Leitão et al., 2015; Battelli et al., 2009). The attention network is often divided into the dorsal and the ventral attention sub-networks (DAN and VAN, or DVAN; Corbetta & Shuman, 2002; Fox et al., 2006). However, we are interested in the whole network due to the continuous interplay between networks, which enables the dynamic control of the different attentional processes (Vossel, Geng and Fink, 2014; Macaluso and Driver, 2005; Sani et al., 2019). We incorporated an active control, the middle temporal area (hMT+), to determine if attention network modulation was specific to parietal stimulation. We also sought to understand whether neural modulation following tRNS coupled with training would lead to long lasting behavioral improvements, and we tested it twenty-four hours following the last training session by examining if behavioral and neuromodulatory changes persisted. Lasting neural and behavioral change would suggest cortical plasticity that outlasts the end of stimulation.

## Materials and Methods

### Regulatory approval

The study was approved by the ethical committee of the University of Trento.

### Participants

Thirty-seven neurologically healthy subjects (20 females, mean age 22.8 years old), with normal or corrected-to-normal vision participated in the study. All subjects were screened for medical contra-indications to MRI and brain stimulation and received monetary compensation for their participation in the experiment.

### Study Design

Participants were randomly assigned to one of three stimulation conditions: (1) bilateral tRNS over parietal cortex (Parietal group), (2) bilateral tRNS over middle temporal cortex, hMT+ area (hMT+ group), and (3) sham stimulation (Sham group). Seven subjects were excluded from the analysis due to head motion during one or both scanning procedures or due to inadequate behavioral performance (i.e. performance was at chance in catch trials), leaving 30 participants for data analysis.

Subjects participated in a multi-session experiment that lasted seven days (one session per day; Figure 1A). On day one, they completed a staircase procedure to measure their psychophysical thresholds for two separate attention tasks: a temporal order judgment (TOJ) and an orientation discrimination (OD) task (described below). On day two (pre-test session), resting-state and the attention task performance measures were taken in the MRI, with trials (from both tasks) randomly interleaved within each block. Participants completed a total of five blocks per task (see section Neuroimaging Procedure for details). On days three to six (training and stimulation sessions), subjects received 25 minutes hf-tRNS or sham while training on the same TOJ and OD tasks, with the two trial types randomly interleaved (cross-tasks training), as in the pre-test (5 blocks per day). Day seven (post-test session) was a repeat of day two.

**Figure 1.**
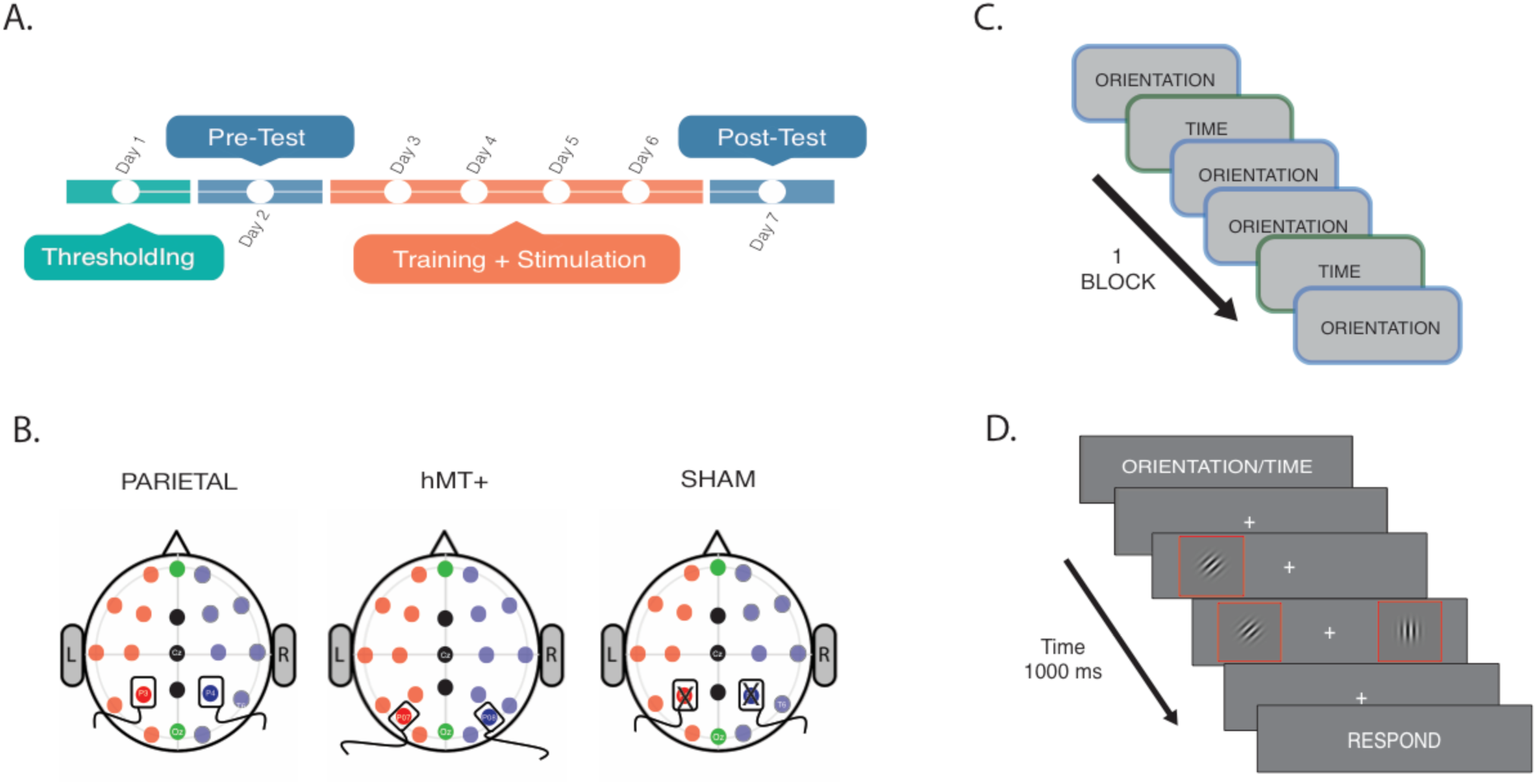
Procedure and stimulation sites. **A)** Experiment Timeline. Thresholds for OD and TOJ tasks were performed on Day 1. On Day 2 subjects were tested on the tasks while fMRI data were collected (pre-test session). From Day 3 to Day 6, subjects underwent behavioral training concurrently with tRNS for 25 minutes. Day 7 was a repeat of Day 1 (post-test session). **B)** Stimulation settings. Location of stimulation sites were localized using EEG 10/20 system. Saline soaked electrodes were placed over P3 and P4 for bilateral parietal and sham stimulation, and over P07 and P08 for hMT+ stimulation. **C)** Intermixed trials sequence example: two types of attention tasks were randomly presented within each block, the OD and TOJ task. **D)**. Example trial. The visual information was the same for TOJ and OD tasks. The cue at the beginning of the trial dictated which feature (time or orientation) should be attended. Red boxes only present for highlighting Gabors in the illustration, and were not present during experimental stimulation.

### Stimuli

Stimuli consisted of a pair of sine-wave gratings (Gabors) positioned 4° to the left and right of a central fixation cross. The Gabors were presented with a spatial frequency of 3 cycles/deg and reduced 50% contrast against a uniform gray background (47.5 cd/m^2^), with a fixation cross positioned in the center of the screen for the entire duration of the trial. The two Gabors were presented with temporal offset asynchronies (for the TOJ task) that corresponded to the individual temporal threshold described in detail below. In addition to the temporal offset, one of the two discs was tilted along the vertical line with a degree that corresponded to the individual orientation threshold calculated for the OD task at baseline. In the TOJ task participants were asked to judge whether two Gabors were presented simultaneously or not, while in the OD task participants were asked to determine whether the two Gabors had the same or different orientation. Throughout the experiment, subjects always viewed stimuli at their individual thresholds (except during catch trials where stimuli were presented above threshold) for both dimensions (time and orientation); however, on any given trial, they had to attend to only one of the features, as instructed by the pre-cue word. The target feature (or “test Gabor”) was presented for equal number of trials (120 trials per trial type, 240 in total per session) on the left and right visual field, within each block per task. Easy (above threshold) catch-trials were also interleaved randomly within each block (n=6, 24 catch trials per session) to check that participants remained engaged and were performing the cross-attention task correctly. In the Easy trials, the two Gabors were presented with large offset asynchronies (≈150 msec, for the TOJ condition), or with a large orientation difference (≈20^°^, for the OD condition). Importantly, in both tasks visual stimuli were presented with temporal offsets (TOJ) and orientation values (OD) fixed at individual threshold levels, which were calculated on day 1.

During days three to six (training days), stimuli were displayed on a 22-in. LCD monitor with a 60 Hz refresh rate controlled by a DELL computer equipped with Matlab r2016a (The MathWorks, Natick, MA) and Psychtoolbox 3.0.8 (Brainard, 1997; Pelli, 1997). Participants were seated 57 cm from the screen in a dark and quiet room, and used a chin rest to ensure consistent positioning. During the fMRI sessions, stimuli were displayed on a Nordic NeuroLab LCD monitor (Basic monitor specs include: 878 mm horizontal x 485 mm vertical; 3840 × 2160 pixels) connected to a Windows PC equipped with the Matlab r2016a and Psychtoolbox 3.0.8, and stimuli were back projected through a mirror inside the scanner.

### Apparatus and Procedure

On day one (Figure 1A), subjects performed two two-alternative forced choice tasks within each block: (1) a temporal order judgment (TOJ) or (2) an orientation discrimination task (OD). Two 3-1 staircase procedures were used to assess thresholds values for the two tasks (TOJ and OD), separately. A single incorrect response decreased task difficulty, while three consecutive correct responses increased task difficulty. The staircase terminated after 30 reversals of the staircase parameter (an average of 60 trials per subject). Temporal offsets and orientation values in remaining sessions were calibrated based on the thresholds to yield 50% accuracy in the same/different judgments. The threshold corresponded to the point of subjective equality (PSE), which represents the threshold value at which the observer experiences two stimuli as identical. The estimated threshold was then used for the last 15 trials and it was considered accurate if subjects performed at 50% on these trials, the expected performance. 50% accuracy was chosen to prevent ceiling effects, thus allowing perceptual learning to occur.

On days two through seven, each trial started with a 2-second instruction interval during which a cue-word indicated which of the two tasks the subjects should perform in the upcoming trial: “Time” instructed the subject to perform the TOJ task (120 trials), and “Orientation” instructed the subject to perform the OD task (120 trials). After the pre-cue, stimuli were presented for 500 msec. Subjects were then prompted to make a forced-choice judgment to indicate whether the two stimuli had same/different orientation (OD task) or if they were presented at the same/different time (TOJ task), depending on the task. During the response interval (1.5 sec), a cue-word was presented to remind subjects which judgment they had to report. A fixation interval of 2 or 4 seconds terminated the trial (trial example, Figure 1D).

### Stimulation Protocol

Before testing, all participants were provided with a short introduction to brain stimulation and safety information. After each participant was briefed, they completed a stimulation safety questionnaire and signed the informed consent. At this time, any participant deemed ineligible for stimulation or MRI procedures was excluded from the experiment. A battery-driven stimulator (DC-Stimulator, NeuroConn, Ilmenau, Germany) was used for electrical stimulation. For the two active tRNS conditions (Parietal and hMT+), 2mA current was applied for 25 consecutive minutes with random alternating frequency delivered at a high frequency range between 101 and 640 Hz. Stimulation was delivered with a fade in/out period of 20 sec at the beginning and at the end of each stimulation session. For sham stimulation, the machine was turned off after the fade-in phase. Two rubber electrodes (size=5⨯7 cm), contained in sponges soaked in saline solution, were placed on the subject’s head and were kept fixed on the stimulation sites with a rubber band and a head cap. Stimulation sites were identified using the International electroencephalographic 10/20 system for scalp electrode localization. The center of the electrode was placed bilaterally over PO7/PO8 (left and right, respectively) for hMT+ and bilaterally over P3/P4 for parietal and sham conditions (Figure 1B). Participants reported no noticeable sensation resulting from tRNS (see also Ambrus, Paulus & Antal, 2010).

### Neuroimaging Procedure

Whole-brain scanning was performed with a 4T Bruker MedSpec MRI scanner using an 8-channel head-coil at the Center for Mind and Brain Sciences of the University of Trento, Italy. High-resolution T1-weighted images were acquired for each subject (MP-RAGE 1×1×1 mm voxel size, 176 sagittal slices). Functional images (T2*-weighted EPIs, TR=2.0 seconds, TE=28.0, flip angle=73°, 3×3×3mm voxel size, 0.99 mm gap, 30 axial slices acquired interleaved, 192 mm FOV) were collected, for a total of 120 volumes per resting state functional run. The event-related task-based neuroimaging scans were then repeated five times. The entire session consisted of one anatomical run followed by six functional runs (one resting-state run, five task runs) and lasted for approximately one hour in total.

During MRI data collection, subjects viewed the stimuli through a periscope mirror mounted on the MR head-coil that allowed the subject to view a screen positioned at the head of the scanner. Throughout the experiment, participants were instructed to maintain fixation on a constantly present central fixation cross. Participants were instructed to indicate the response with an MR compatible response box (two double-buttons response pads) with left index finger indicating “same” and right index finger indicating “different” responses.

### Data Analysis

#### Behavioral data analysis

Statistical analyses were performed using MATLAB (The MathWorks) and SPSS. We only present and compare behavioral data from day 2 and day 7 (pre and post-test days), the two sessions during which fMRI data were also collected, thus data from the training (day 3 to 6) will not be reported. Accuracy scores were calculated as a proportion of correct responses from the recorded responses on each day. We analyzed data separately for the two tasks (OD and TOJ) and we compared pre-post stimulation performances. For both pre-and post-stimulation, we first checked for normality and homogeneity using Levene’s test. Next, we performed two mixed repeated measures ANOVA to determine the effect of stimulation condition on performance in the two attention tasks across sessions. The within-subjects factor was Day (n=2 for pre-post stimulation analysis) and the between-subjects factor was Stimulation condition (n=3). For the post-pre-stimulation analysis, we first tested whether the three stimulation groups differed at baseline, then calculated the total improvement as the difference between the post and pre-stimulation performance, thus obtaining the delta value for each stimulation condition (ΔPerformance|Improvement=PostStimulation - Performance–PreStimulation Performance). We next performed a one-way ANOVA to test whether there was a significant difference between the performance improvements (delta values) between stimulation conditions.

### Resting State fMRI data analysis

Preprocessing of resting-state functional scans was initially performed using BrainVoyagerQX (Brain Innovations Inc., Maastricht, The Netherlands) and included the elimination of the first 4 volumes due to non-steady state magnetization. Images were subsequently subjected to 3D motion correction, slice-timing correction, realignment, and spatial smoothing with a 6 mm FWHM Gaussian kernel. Runs with instantaneous head motion exceeding 3mm in any given dimension were discarded. The functional scans were manually co-registered to each individual’s high-resolution anatomical scan, then normalized into standardized Talairach coordinates (Talairach & Tournoux, 1989).

Advanced pre-processing of the functional data from targeted regions of interest (ROIs) was performed in MATLAB (The Math Works) to remove data contamination caused by motion, which can alter functional connectivity scores. This pipeline included nuisance regression using 12 motion estimate regressors (the six head realignment parameters obtained by rigid body head motion correction and their first-order derivatives) and estimates of global signal obtained from the ventricles. The ROI timeseries were then detrended and volumes with movement-related activity were removed (scrubbing) and temporarily replaced with resampled volumes (computed using LSSD Fit) in order to apply band-pass filtering (0.009-0.08Hz; Cordes et al., 2001). The censored and resampled volumes were then removed from further analysis. After these processing steps were applied, only those runs that retained at least 3 min worth of data were used for further analyses (Satterthwaite et al., 2013).

We computed functional connectivity on resting-state data using 10 bilateral regions of interest (ROIs): the intraparietal sulcus (IPS), human middle temporal complex (hMT+), and frontal eye fields (FEF) of the dorsal network; and the temporal parietal junction (TPJ) and ventral frontal cortex (VFC) of the ventral network. These regions were identified using standardized mean coordinates (Table 1) derived from the literature (Battelli, Grossman, & Plow, 2017; Vossel, Geng and Fink, 2014; Corbetta et al., 2008; Fox et al., 2006). Functional data from all voxels within a 6mm radius sphere from the center coordinate were averaged into a single timeseries, keeping the ROIs similar in size across participants. The time courses were extracted from each ROI in the resting-state run and functional connectivity scores were computed as the Pearson’s r correlation coefficient. Correlation coefficients were Fisher-z transformed prior to subsequent statistical analyses.

**Table 1.** Brain Coordinates. Group mean Talairach X, Y and Z coordinates for the centroid of each region of interest (Left and Right hemisphere) for the DVAN.

| ROIs Standardized Coordinates |  |  |  |  |  |  |  |
| --- | --- | --- | --- | --- | --- | --- | --- |
|  |  | LEFT HEMISPHERE |  |  | RIGHT HEMISPHERE |  |  |
|  |  | X | Y | Z | X | Y | Z |
| DVAN | IPS | -26 | -63 | 48 | 25 | -60 | 48 |
|  | hMT+ | -44 | -67 | 3 | 44 | -71 | 6 |
|  | FEF | -30 | -5 | 48 | 30 | -5 | 48 |
|  | TPJ | -48 | -54 | 26 | 48 | -54 | 21 |
|  | VFC | -42 | 12 | -1 | 40 | 17 | 4 |

To test the effect of training and stimulation site, functional connectivity scores were subjected to a repeated measures Analysis of Variance with the within-subject factor session (n=2) and between-subject factor of stimulation site (n=3). The difference between functional connectivity scores in the first and last MRI sessions was performed on individual subjects to evaluate the delta in FC scores pre- and post-stimulation (Δ rs-FC = rs-FC(S2) – rs-FC(S1)). Post-hoc pairwise comparisons of the delta FC (t-test) were next performed between stimulation conditions. Finally, to explore the relationship between functional connectivity within the DVAN and behavioral performance, we computed the linear regression between each participant using individual measures of behavioral improvement (accuracy changes pre-post stimulation) against individual measures of stimulation-induced changes in rs-FC. Using Pearson correlation coefficient, we hence calculated the correlation between behavioral changes and brain activity modulation.

## Results

### Learning-dependent changes in behavior

#### Orientation Discrimination

We first ensured that the three stimulation groups did not differ prior to stimulation by comparing their behavioral performance on the orientation task. A one-way ANOVA revealed no significant difference between groups prior to stimulation (F(2, 27)=.575, p=.569). We then examined the combination of stimulation and training on learning. This analysis revealed a main effect of session on orientation discrimination performance (F(1, 27)=10.056, p=.004), indicating that accuracy scores changed as a function of training. There was no significant main effect of stimulation condition alone on performance (F(2, 27)=.273, p=.763). Crucially, the analysis revealed a significant interaction between stimulation condition and session (F(2, 27)=11.257, p<.001), indicating that learning from the pre-to post-stimulation sessions varied significantly depending on the stimulation site.

To further investigate the significant interaction effect, we ran post-hoc pairwise t-tests to compare pre-post stimulation performance for each group, separately. As depicted in Figure 2A, we found that only the Parietal group showed significant improvement in orientation judgment accuracy between the pre- and post-stimulation sessions (t(9)=-4.53, p=.001, average improvement 29.8%), while there were no significant differences between pre-post stimulation performance for the hMT+ group (t(9)=-.608, p=.558, average improvement 2.2%), nor for the Sham group (t(9)=.602, p=.562, average decrease −3.1%). A one-way ANOVA on performance improvements values confirmed a significant difference between the three stimulation groups (F(2, 22.57)=11.257, p<.001, Brown-Forsythe corrected F value). Bonferroni corrected post-hoc pairwise comparisons revealed a significant difference in performance between Parietal and Sham condition (p=.003), and between Parietal and hMT+ (p=.001), while hMT+ and Sham did not differ significantly (p=.832).

**Figure 2.**
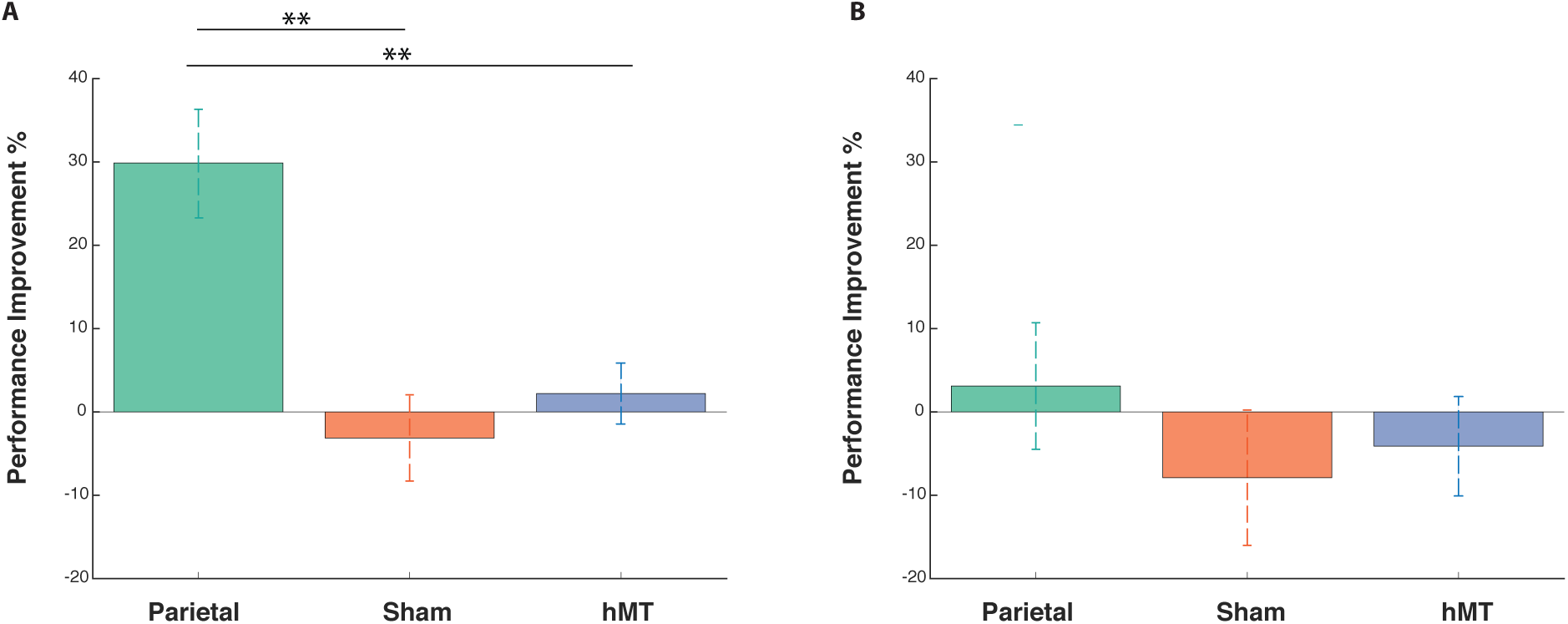
Behavioral change on the OD and the TOJ task following stimulation and training. (A) Improvement for all conditions on the OD and (B) the TOJ task normalized relative to performance at baseline prior to stimulation. Colored bars indicate the percentage accuracy difference (delta value) between pre- and post-stimulation session per group: green, orange and blue for parietal, sham and hMT+ group, respectively. Asterisks indicate significant difference.

#### Temporal Order Judgment

A one-way ANOVA on pre-stimulation session data revealed no significant differences between the groups at baseline (F(2, 27)=0.14, p=.867). We next investigate whether there was a significant difference between the pre- and post-stimulation TOJ performance depending on stimulation condition. A mixed factors repeated measures ANOVA revealed no main effect of session (F(1, 27)=.496, p=.48) nor of condition (F(1, 27)=.669 p=.521) on performance. There was also no significant interaction between session and stimulation condition (F(2, 27)=589, p=.562), which indicates that performance in the pre and post-stimulation session did not differ depending on stimulation condition.

A one-way ANOVA on calculated delta values (difference between pre- and post-stimulation performance) confirmed a no significant performance change between stimulation conditions (F(2,27)=.589, p=.562). Participants in the parietal tRNS group performed 3.1% better following multi-session stimulation and training, while there was a slight decrease in performance for the hMT+ (−4.1%) and the sham group (−7.8%), as depicted in Figure 2B.

### Stimulation and learning-dependent changes in functional connectivity

We next examined the effect of multi-session hf-tRNS and training on resting state functional connectivity (rs-FC) of the Dorsal and Ventral Attention Network (DVAN). First, we calculated the mean FC per subject within the DVAN by averaging the FC scores of the 45 ROI pairs, we then averaged across subjects in each group (Van Den Huevel et al, 2017; Nicolini et al., 2019). We compared FC prior to stimulation between groups and found no significant difference (F(2, 27)=.3, p=.743), indicating that FC did not differ between stimulation groups at baseline (pre-stimulation session).

A mixed repeated measures ANOVA testing the two factors of session (within-subjects factor, n = 2) and stimulation site (between-subjects factor, n = 3) on functional connectivity scores revealed no main effect of training on FC scores (F(1, 27)=.001, p=.986) and a main effect of stimulation site (F(2, 27)=4.064, p=.029), indicating that the connectivity among the DVAN nodes changed as a function of where tRNS was applied. Importantly, the interaction between stimulation condition and session was found to be highly significant (F(2, 27)=11.507, p <.001), indicating that the change in rs-FC scores differed depending on the stimulation group (Figure 3) and this change was persistent the day after the end of the combined training protocol.

**Figure 3.**
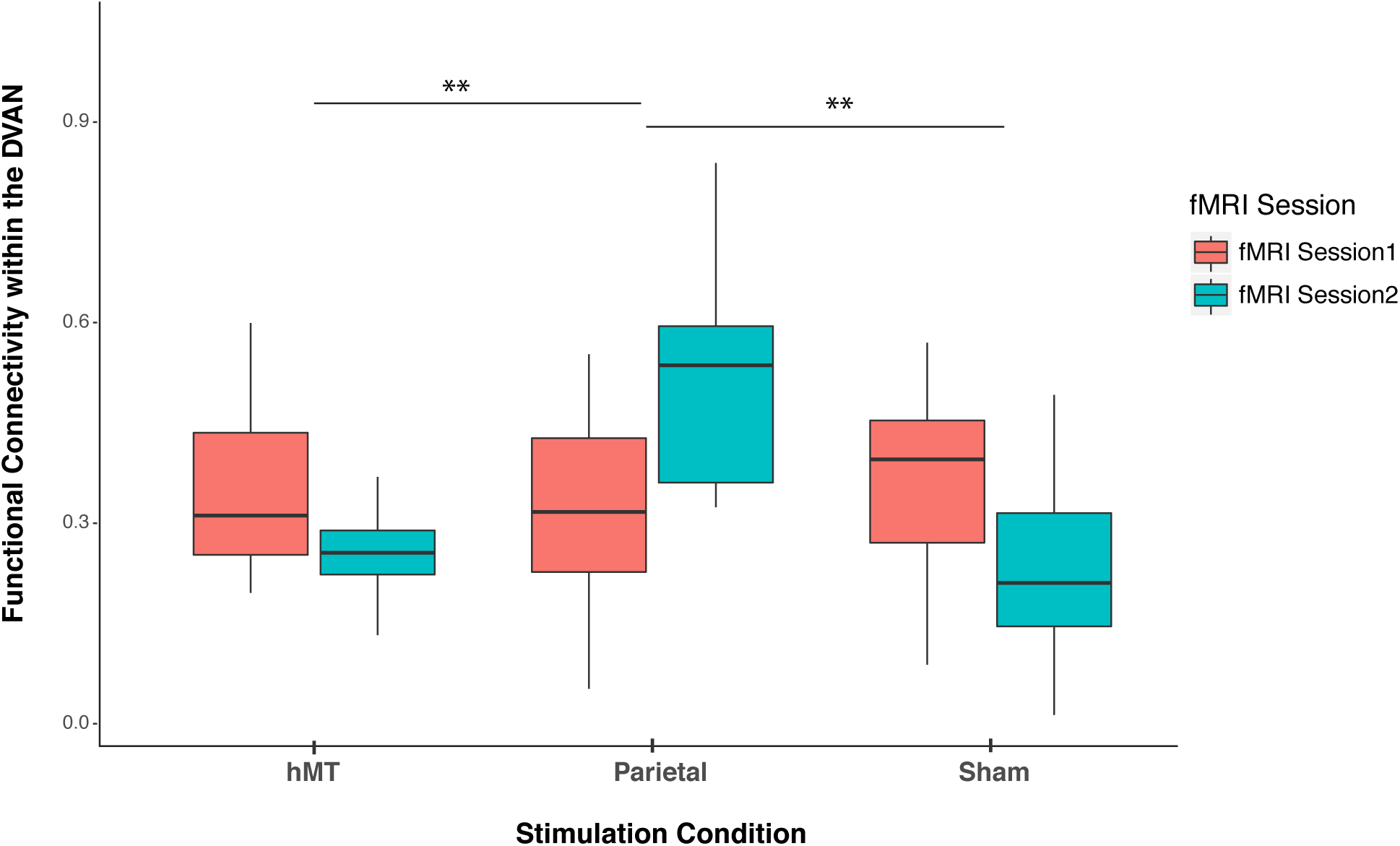
Pre- and post-stimulation FC changes. Overall mean Functional Connectivity of the Dorsal and Ventral Attention Network averaged per stimulation condition (Parietal, hMT+ and Sham) and per fMRI session (pre- and post-stimulation session). Asterisks indicate significant difference.

To detect whether FC was impacted by stimulation condition on each day, we compared the rs-FC of the three stimulation groups at each time point (pre- and post-stimulation day) by performing two one-way ANOVA analyses, one per session. One-way ANOVA analysis on post-stimulation session revealed a significant effect of stimulation condition on FC scores (F(2, 27)= 11.677, p<.001), which indicates that FC differs on this day between stimulation conditions (Figure 3). We then ran t-test comparisons on post-stimulation FC scores to reveal simple effects between pairs of stimulation conditions. We found that FC scores were significantly different in the Parietal group compared to the hMT+ group (t(18)=3.863, p=.002) and compared to those of the Sham group (t(18)=3.64, p=.001), while FC did not differ significantly between Sham and hMT+ groups (t(18)=.187, p=.854), (Figure 3). We further investigated the effect of stimulation on FC by comparing pre and post-stimulation FC values within each stimulation condition. Pairwise t-test comparisons revealed that FC scores were significantly different between the pre- and post-stimulation session in the Parietal (t(9)=-2.973, p=.016) and the hMT+ (t(9)=-2.99, p=.015), while they were no significant in the Sham group (t(9)=2.217, p=.054). Resting state functional connectivity patterns increased within the main nodes of the DVAN after parietal stimulation only, while it decreased after hMT+ and Sham stimulation (Figure 3). Post-stimulation averaged correlation values are depicted per condition in Figure 4.

**Figure 4.**
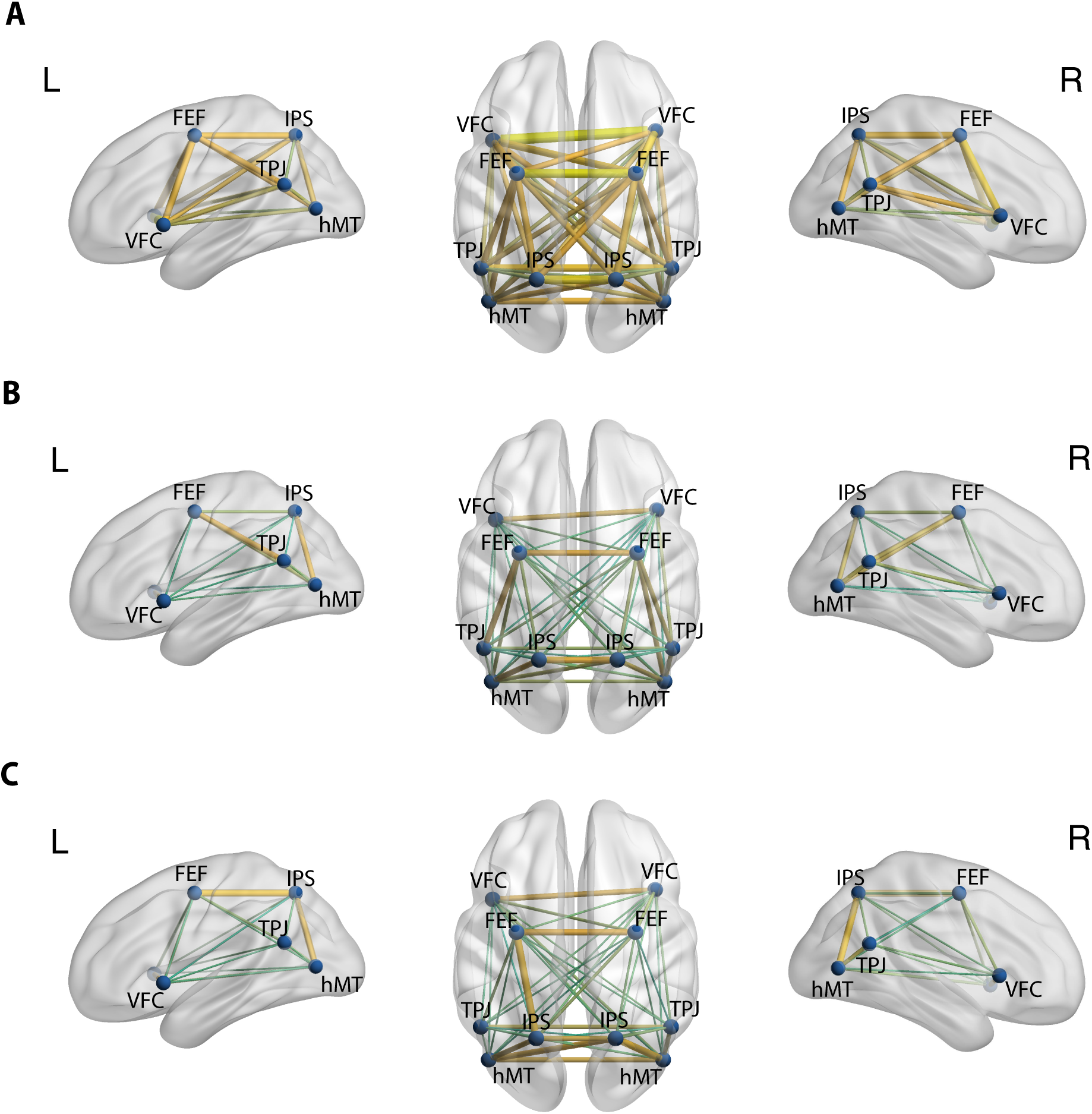
Post-stimulation correlations between pairs of ROIs are projected onto inflated brain representations. Color and thickness of the edges between nodes indicate the strength of correlation for Parietal (A), hMT+ (B) and Sham (C).

In a second analysis, we investigated the modulation of the DVAN by taking into consideration all ROIs pairs as single components of the network per subject (n=45 ROIs to ROIs pairs per subject), instead of using one averaged correlation value for the whole network as in the previous analysis. We performed a mixed repeated measures ANOVA to test the effect of the stimulation site and session on all correlation values within the DVAN. This analysis revealed no main effect of session on FC scores (F(1, 1347)=.002, p=.964), but a main effect of stimulation group (F(1, 1347)=23.07, p=<.001). Importantly, the interaction between stimulation group and session was found to be strongly significant (F(2, 1347)=80.206, p<.001), which indicates a significant difference between rs-FC scores on the two sessions depending on the stimulation condition (Figure 5 and Figure 6).

**Figure 5.**
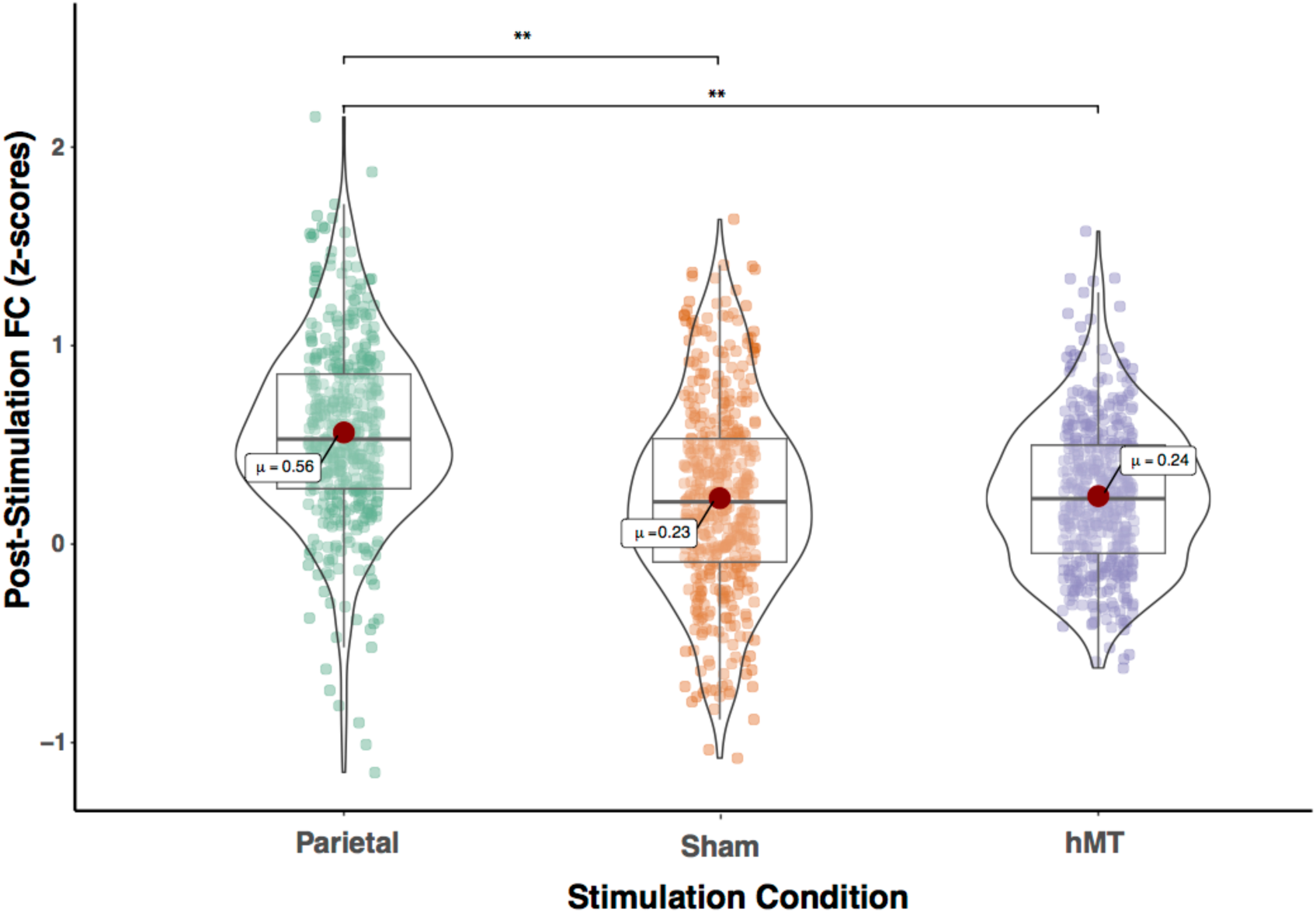
Functional connectivity within the DVAN after multi-session hf-tRNS coupled with training (post-stimulation session). Correlations of all ROI pairs (z-scores) constituting the DVAN are represented separately for the three stimulation conditions (Parietal, Sham and hMT+). Asterisks represents significant differences (**p<0.001).

**Figure 6.**
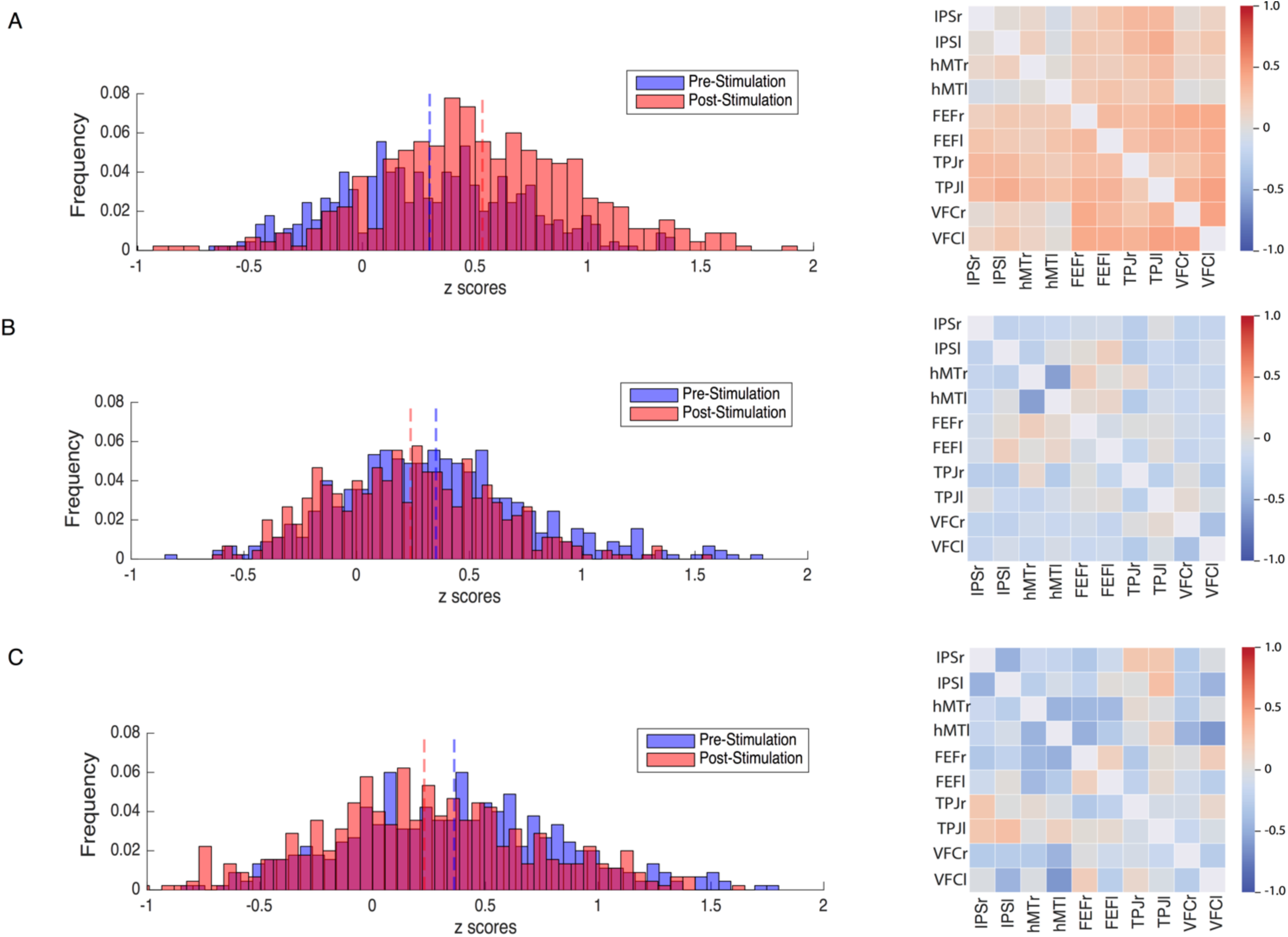
Modulation of Resting-state Functional Connectivity of the DVAN. **Left Panel:** z-scores frequency distribution pre-(light blue bars) and post-(light red bars) multi-session hf-tRNS coupled with training per each stimulation condition (A. Parietal, B. hMT+, C. Sham). Dark red indicates the overlapping distribution between pre- and post-stimulation. Dotted red and blue lines indicated the mean for each z-scores distribution. **Right Panel**: Correlation matrices represent the result of the computed difference between FC scores pre- and post-stimulation (Δ=FC(S2) - FC(S1)) depicted by condition (A. Parietal, B. hMT+, C. Sham). Each correlation difference between ROI pairs was first calculated at subject level and then averaged across subjects. Colors bars indicate the strength and direction of correlation change values for each regions’ pair (red colors indicate higher connectivity; blue colors indicate lower connectivity).

We then compared the rs-FC on all single components (pairs) of the post-stimulation session by performing a one-way ANOVA. Because Levene’s test indicated that the assumption of homogeneity of variances was violated, the Brown-Forsythe F-ratio is reported. This analysis revealed a strong significant effect of stimulation condition on post-stimulation FC scores (F(2, 1286.64)= 81.839, p≤ .001), which indicates that FC was significantly different on this session among the three stimulation condition. T-test comparisons on post-stimulation FC scores (Figure 5) revealed that connectivity was significantly increased in the Parietal group compared to the hMT+ (t(865.23)=11.59, p=<.001, df corrected for inhomogeneity of variance) and to the Sham group (t(892.6)=10.518, p<.001, df corrected for inhomogeneity of variance). Post-stimulation FC scores were not significantly different between the hMT+ and the Sham group (t(837.68)=.357, p=.721, df corrected for inhomogeneity of variance).

Figure 6 shows how connections between the nodes of the DVAN changed after Parietal stimulation resulting in stronger positive correlations within the network nodes, while the connections between the nodes after hMT+ and Sham stimulation resulted in a more negative correlation, indicating decreased connectivity.

### Functional connectivity-behavior correlation

Next, we examined the relationship between changes in behavioral performance and the resting-state FC within the DVAN. We found subjects that displayed a higher performance also tended to show higher rs-FC within the DVAN (measured as the mean of the correlations within the nodes of DVAN) in the post-stimulation session (r=0.40). The linear regression model revealed a significant regression coefficient (p=0.019, R-squared=0.163) which indicates that changes in functional connectivity are significantly associated with changes in behavioral scores in the final session (Figure 7).

**Figure 7.**
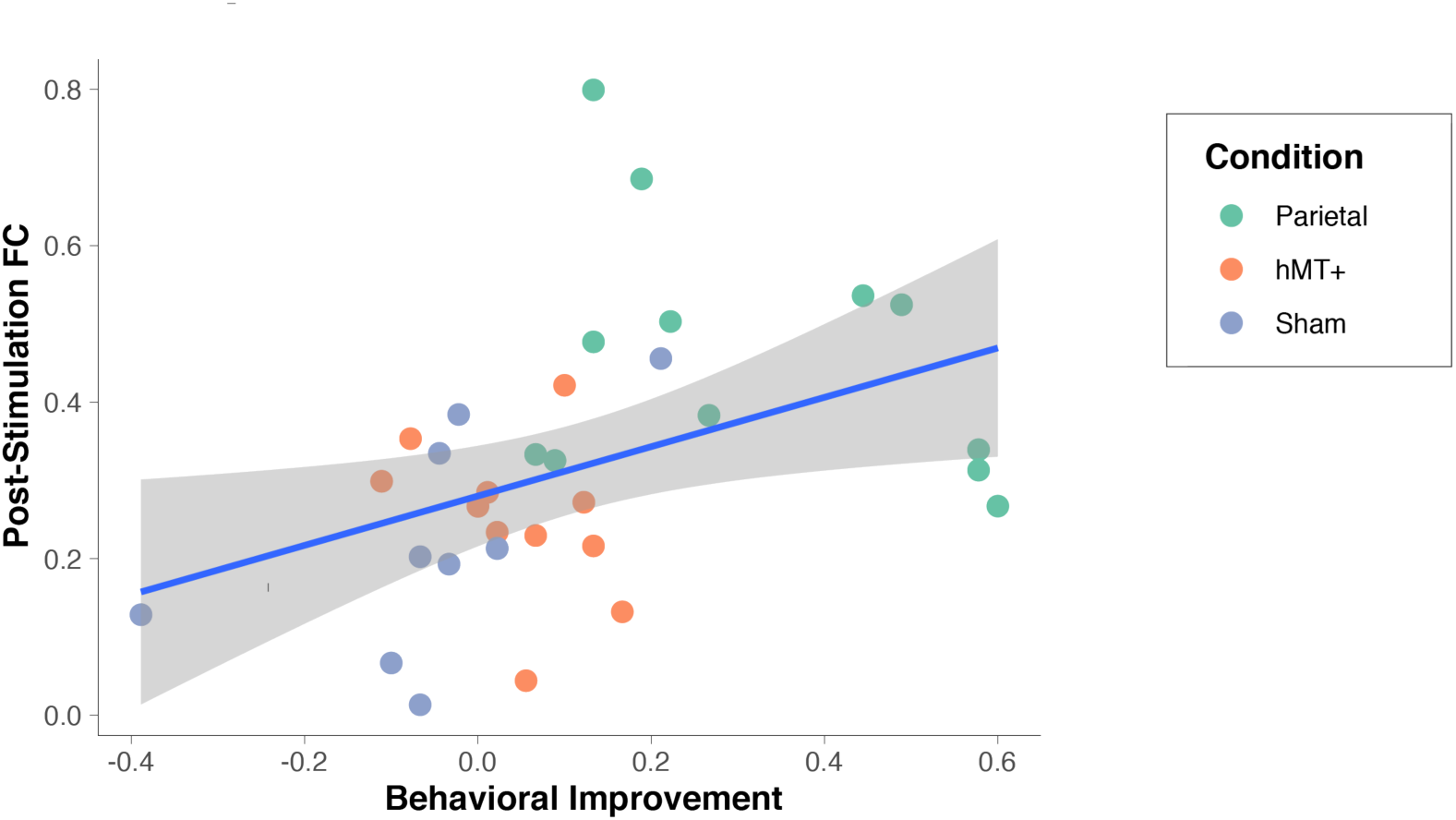
Changes in Resting-State Functional Connectivity correlate with changes in behavioral performance. Scatter plot showing the positive correlation between post-stimulation rs-FC values within the Dorsal and Ventral Attention Network (DVAN, y-axis) with behavioral improvement on the Orientation Discrimination Task (Behavioral difference between pre and post-stimulation sessions, x-axis).

## Discussion

In this study we investigated the potential for visual training coupled with noninvasive brain stimulation to promote significant improvements in behavior and increase cortical functional connectivity. We trained subjects on cross-tasks training to optimize learning in an attention-based learning paradigm over multiple days. We combined this training with high frequency transcranial random noise stimulation (tRNS) using a protocol optimized to facilitate cortical plasticity with the goal of increasing and speeding up training improvements relative to training without tRNS. Our stimulation protocol, when applied over parietal cortex, dramatically shortened the learning period, overcoming the limitations of previous training paradigms. Crucially, tRNS coupled with training also increased functional connectivity within the attention cortical network when delivered over parietal cortex, and the connectivity pattern changed more strongly for individuals with the largest behavioral improvement after training. Thus, we conclude that tRNS has potential to strengthen task-relevant connections between sensory and attention systems to promote learning and plasticity.

Bilateral tRNS over the parietal cortex during training resulted in a strong boost in performance, while tRNS over hMT+ and sham had no effect on behavior. Crucially, improvement in behavior in the parietal condition positively correlated with increase in functional connectivity within the main nodes of the dorsal and ventral attention networks, indicating that concurrent training and stimulation modulated large-scale cortical dynamics, strengthening functionally selective neural pathways (Chiappini et al., 2018), a potential physiological marker of enduring changes (Polania et al., 2011).

We also sought to understand whether a significant improvement could be achieved with a short training protocol, in conditions where we normally would not expect learning to occur (Huang et al., 2012; Aberg et al., 2009; Li et al. 2004). We adopted a cross-task training that uses the same visual stimulus for two different tasks (Szpiro et al., 2014), and we coupled the training with a stimulation protocol during the learning acquisition phase. Results indicate that it is the combination of a challenging task with concurrent brain stimulation over the relevant cortical circuits that facilitates fast learning, a paradigm amenable to promote plasticity.

Together, these behavioral results suggest that learning of a visuospatial attention task was achieved within few training sessions and that this facilitation was long-lasting, as increased performance outlasted the end of stimulation for at least 24 hours, when the last fMRI session after the end of training was performed. These findings extend the results of a previous experiment that tested tRNS efficacy upon attention within one session, but found a rapidly decaying beneficial effect upon the end of stimulation (Tyler et al., 2018).

However, using a multi-session stimulation approach coupled with training promoted retention of the beneficial effect of training, as also shown by the results in increased resting state functional connectivity after training (Chen et al., 2015; Sarabi et al., 2018; Kang et al., 2018).

There might be several conditions that facilitate the emergence of learning, all likely driving the stimulus related activity to reach the learning threshold (Seitz & Dinse, 2007). First, in this study we demonstrate that stimulation site selection within a network is crucial to obtaining successful augmentation. Performance improved only when tRNS was delivered to the parietal cortex, and was absent in all other conditions, including stimulation to hMT+, often included as a node of the dorsal attention network. Thus, our results indicate the parietal lobes have a direct involvement in visual perceptual learning and attention processes (Giovanelli et al., 2010, Law and Gold, 2008; Shadlen and Newsome, 2001; Colby and Goldberg, 1999). Second, these results indicate that tRNS effects do not spread to regions in close anatomical proximity, but rather spread within strongly functionally connected circuits. Stimulation to hMT+, which is close to the parietal lobes, did not have any effects on behavior nor on FC. Third, our findings support the idea that learning is facilitated when the cortical processes underlying the task are stimulated while in an aroused state derived by task performance (Wright et al., 2010), and indicate the possibility to use tRNS to rapidly reach this optimal state. Importantly, our study further indicates the benefits of short but effective training sessions, an ideal condition to promote learning (Censor and Sagi, 2008; Censor et al., 2016).

One might ask why we observed learning occurring for one attention task only. Our results are consistent with previous studies that used a similar cross-task training paradigm and found behavioral improvement in only one of two tasks trained (Szpiro, Wright and Carrasco, 2014; Wright et al., 2010). One potential explanation for task selective improvement is that making two different perceptual judgments on the same visual stimulus may require tuning of different cortical representations (Li et al., 2004). Simultaneous training might induce competition between these two representations, ultimately leading to the enhancement of one and inhibition of the other, as suggested in the augment Hebbian reweighting model (AHRW; Dosher et al., 2013; Dosher & Lu, 2017). Similar to Szpiro and colleagues’ study, when our participants trained on two randomly interleaved tasks using identical visual stimuli, improvements were observed only in the orientation discriminations, suggesting the training may have changed perceptual weighting of the orientation channels.

In line with this hypothesis, our study showed that orientation PL was apparent with our very short training sessions only when coupled with tRNS over parietal cortex, further emphasizing the role of neuromodulation in speeding up the learning process. In the parietal stimulation group, we positioned the electrodes over the EEG locations P3 and P4, roughly corresponding to the posterior intraparietal sulcus. The IPS has been linked to visuospatial attention in several imaging and brain stimulation studies (Corbetta and Shulman, 2011, Battelli et al., 2009; 2017), and the orientation discrimination task involves the processing of visuospatial components. Spatial processing was not relevant in the temporal order judgment condition and that is typically associated to more ventral areas (Chica et al., 2011; Agosta et al., 2017; Husain & Nachev, 2007). The temporoparietal junction (TPJ), at the crossroad between the ventral and the dorsal visual pathways has been linked to temporal attention (Tyler et al., 2015: Battelli et al, 2007; Husain and Rorden, 2003). Although this explanation is rather speculative, the selectivity of the behavioral effect along with functional connectivity results might indicate that the IPS plays a pivotal role in orientation discrimination across the visual space (Capotosto et al., 2013; Corbetta & Shulman, 2002). Moreover, the parietal lobe has been associated with selective and gating attention mechanisms (Suzuki & Gottlieb, 2013), and tRNS might have facilitated these gating mechanisms during learning.

Finally, some studies have analyzed the importance of task engagement or difficulty during stimulation, and have demonstrated how these factors affect the behavioral outcome (Bortoletto et al., 2015; Hsu, Juan and Tseng, 2016). In our study, participant engagement was challenged in two ways. First, task difficulty was individually tailored and equated for difficulty across subjects ensuring subjects’ engagement with the task (Waskom et al., 2019). Second, the experimental design randomly alternated between experimental tasks, forcing the continuous switch of attention between stimulus features, might force a strategy that favors one task due to limited available cognitive resources.

We next asked whether there were cortical physiological changes that indicate more sustained plastic changes along with behavioral change, and if such changes were within task-related well established attention-related cortical networks. Specifically, we analyzed whether the multi-session tRNS had an effect on functional connectivity at rest in the dorsal and ventral attention network (DVAN), which we targeted through stimulation. Brain regions in this network mediate attention mechanisms and consequently shape the analysis of visual perceptual input (Kastner & Ungerleider, 2000). The dorsal/ventral attention network plays a pivotal role in attention functions and constitutes a large occipital-frontoparietal network that controls allocation of attention to positions and visual features in space (Kastner & Ungerleider, 2000; for a review see Fiebelkorn & Kastner, 2019: Vossel, Geng and Fink, 2014). Tasks requiring selective attention have been shown to activate the ventral part of the attention network (VAN; Corbetta et al., 2008; Vossel et al., 2012), while the dorsal portion of the attention network, in particular the IPS and the FEF regions, is active during many visuospatial tasks that require spatial updating and the selection of important regions of space (Kastner and Ungerleider, 2000; Corbetta et al., 1998; Merriam, Genovese & Colbi, 2003; Greenberg et al., 2010).

Consistent with the relevance of both ventral and dorsal streams of the attention network for the performance of the task used in this experiment, our results show increased functional connectivity within the entire network. Almost all connections between the node pairs displayed higher connectivity after multi-session parietal stimulation only. This result indicates that bilateral hf-tRNS over the parietal lobes modulated the whole attention-related network, instead of producing local changes alone. Importantly, the neural modulation was network specific, not a propagation of stimulation across neighboring cortical regions. This was demonstrated in the hMT+ stimulation group, where the neighboring IPS was not modulated by hMT+ stimulation. Similar to our results, previous physiological studies on non-human primates found changes in the lateral intraparietal area but not in the MT area suggesting that perceptual learning requires the involvement of higher areas that induce changes in how the sensory information is interpreted to drive behavior (Law and Gold, 2008). The lack of improvement in the hMT+ stimulation condition might indicate that while this cortical area might be involved in attentional processing (Yates et al., 2017; Yao et al., 2016), for other functions involved in learning, such as decision processing, the parietal lobe plays a more pivotal role (Herrington and Assad, 2010; Zhou and Freeman, 2019). Our results are also consistent with previous research that found persistent network dynamics changes following other types of stimulation (e.g., transcranial magnetic stimulation, TMS; Battelli et al., 2017; Ruff et al., 2006; 2007; Lee & D’Esposito, 2012), and points to the possibility to use our neuromodulation protocol to exert long-lasting and distal effects.

Functional connectivity changes following parietal stimulation are likely to have affected behavior and driven the system to better respond to the demands of the attention task. Rapidly evolving changes in connectivity between interconnected regions has been proposed as the key mechanism that allows large-scale networks to respond to task demands (Buschman and Kastner, 2015). In the context of tRNS, we speculate that a fast modulation of the network connectivity can be due to the temporal summation of neural activity, as well as to the offline stochastic effect of tRNS (Chaieb et al., 2015). In fact, the stochastic resonance mechanism has been proposed to lead to higher synchrony of oscillations between neurons across large-scale functional systems, which in turn creates strong links between firing neurons (van der Groen & Wenderoth, 2016; van der Groen et al., 2018; Schwarzkopf et al., 2011; McDonnell & Abbott, 2009; McDonnell & Ward, 2011). The strong correlation found between improved performance and increased FC suggests the pivotal role of the IPS in network mediating these system effects during perceptual learning (Szczepanski & Kastner, 2013).

## Conclusions

This multi-method study investigates the potential long-term benefits of tRNS on cortical plasticity and its ability to efficiently alter cortical networks to promote attention and visual perceptual learning. Our neuroimaging and behavioral results support the idea that the modulation of task-related neural networks induced by tRNS can efficiently promote behavioral improvements and speed up the emergence of learning. Overall, this is the first study demonstrating the sustained and selective nature by which tRNS operates on cortical dynamics, and consequently opens a critical window during which the cortex might be more plastic and responsive to reorganization needs to promote learning. Clinical and healthy populations could significantly benefit from the shortened learning time afforded by the combination of tRNS with behavioral learning.

## Acknowledgements

The present study was funded by the Autonomous Province of Trento, Call “Grandi Progetti 2012”, project “Characterizing and improving brain mechanisms of attention – ATTEND” (FC, LB); EG was supported by the National Science Foundation BCS0748314. Any opinions, findings, and conclusions or recommendations expressed in this material are those of the author(s) and do not necessarily reflect the views of the National Science Foundation.

